# Novel repressors of cambium activity in *Arabidopsis*

**DOI:** 10.64898/2026.01.28.702292

**Authors:** Xing Wang, Jingyi Han, Emma K. Turley, Riikka Mäkilä, Anne-Maarit Bågman, Julia M. Kraus, Qing He, Hanan Alhowty, Joanna Edwards, Yuqi Li, Raluca Blasciuc, Wiktoria Fatz, Wenbin Wei, Miguel de Lucas, Siobhán M. Brady, Shixue Zheng, Chunli Chen, Ari Pekka Mäh-önen, J. Peter Etchells

## Abstract

Wood is the greatest reservoir of terrestrial biomass and an essential carbon sink. Formed of xylem, it is derived from the cambium, a meristematic zone within plant stems from which phloem also forms. In Arabidopsis, cell division within the cambium is promoted by three major factors: auxin, cytokinin, and the TDIF-PXY ligand-receptor pair. Meristems and other stem cell populations are typically regulated by a balance between cell division-promoting factors and those that repress cell division to control meristem size, however few factors with cambium-repressing activity are known. Here we combined transcriptomics and transcriptional network analysis, which led to identification of related homeodomain zinc-finger transcription factors, ATHB23, ATHB30, and ATHB34, that repress cambium activity. These factors inhibit cambium activity by directly binding of promoters from a subset of auxin, cytokinin and TDIF-PXY transcriptional target genes, resulting in attenuation of their transcription. Our findings thus reveal a new mechanism underpinning balanced cambium activity.

## Introduction

Stem cell niches are typically regulated by opposing molecular mechanisms that balance pluripotency, cell division, and differentiation. Loss of stem cell promoting mechanisms can result in too few cell divisions and/or consumption of the niche itself as cells prematurely differentiate. An overactive stem cell niche often produces disorganised organs due to failure of differentiation programmes. These features are characteristic of the vascular cambium, the meristem that is responsible driving secondary (radial) growth in the stems of seed plants. Here, a bifacial stem cell population forms xylem and phloem on opposing sides (Shi et al. 2019; Bossinger and Spokevicius 2018; Smetana et al. 2019), with xylem constituting the woody tissue. As such the cambium is the stem cell population responsible for the generation of the majority of terrestrial biomass (Bar-On et al. 2018).

In Arabidopsis, factors that promote cambium activity include cambium-expressed AIL (CAIL) transcription factors (Randall et al. 2015; Eswaran et al. 2024). Their expression defines the cambium stem cells and is promoted by TDIF-PXY (Eswaran et al. 2024), a ligand-receptor pair which acts redundantly with a second signalling mechanism characterised by the receptor kinase, ERECTA (ER)(Fisher and Turner 2007; Hirakawa et al. 2008; Wang et al. 2019). TDIF-PXY additionally activates WOX4 and WOX14, transcription factors that promote cambium cell proliferation(Hirakawa et al. 2010; Etchells et al. 2013). WOX14 forms a coherent feed-forward loop with two further transcription factors TMO6 and LBD4, with WOX14 activating expression of *TMO6* and *LBD4*, and TMO6 also activating expression of *LBD4* (Smit et al. 2020). *TMO6* and *LBD4* are members of multi-gene families that are regulated by cytokinin (Ye et al. 2021; Smet et al. 2019), while auxin also governs the activity of genes within this network (Smetana et al. 2019; Suer et al. 2011). Factors opposing cambium activity include the transcription factor BES1 and HD-Zip III transcription factors (Saito et al. 2018; Baima et al. 2001; Carlsbecker et al. 2010; Smetana et al. 2019). TDIF-PXY marks BES1 for turnover, excluding it from the cambium, which is necessary for cambium activity as BES1 binds to the promoter of *WOX4*, repressing its expression (Kondo et al. 2014; Hu et al. 2022). HD-Zip III’s are known to repress *CAIL* transcription in xylem initial cells (Eswaran et al. 2024). However, the limited number of known repressors of cambium activity leaves an imbalance in the system, which in turn suggests that others are present waiting to be described.

Here, we identified ATHB34 and its homologues ATHB23 and ATHB30 as novel repressors of cambium activity, by cross-referencing transcriptomic data with a transcriptional regulatory network. ATHB34, ATHB23 and ATHB30 bound the promoters of *WOX14, TMO6*, and *LBD4*, repressing their expression and promoting differentiation via direct repression of genes that drive proliferation in the cambium. As the cambium is the meristem from which wood is derived, these results reveal a molecular mechanism by which proliferation and differentiation are balanced in production of this versatile biomaterial.

### Genes regulating cambium development

Higher order mutants between *PXY* and *ER* families are characterised by profound secondary growth defects (Wang et al. 2019). It is likely that genes involved in regulating cambium homeostasis are differentially expressed in these backgrounds, providing an avenue for gene discovery. The secondary growth region of 3-week-old roots from wild type, *pxy pxl1 pxl2* (*px*f), *er erl2*, and *px*f *er erl2* lines were analysed for tissue composition (**Fig. 1a-e and Supplementary Fig. S1a-d**) prior to RNA-seq. Vascular cylinder area and the total number of vascular cells were rduced in *px*f *er erl2* mutants relative to other lines (**Fig. 1a-c**). However, the number of xylem vessels remained unchanged (**Fig. 1d**), meaning these plants had a higher proportion of xylem vessels overall. We also determined the position of the xylem vessels relative to phloem cells. In wild type, xylem and phloem are derived from opposing sides of the cambium. The distance from the primary xylem at the centre of the root to the differentiating xylem vessels is consequently always smaller than the distance to differentiating phloem elements (**Supplementary Fig. S1a**). As such the ratio of these distances is less than 1. We used this ratio as a proxy for xylem and phloem positioning, finding it to be higher in *px*f *er erl2* mutants than in controls (ratio of 1.41 compared to 0.75 in wild type; **Fig. 1e**).

**Figure 1:**
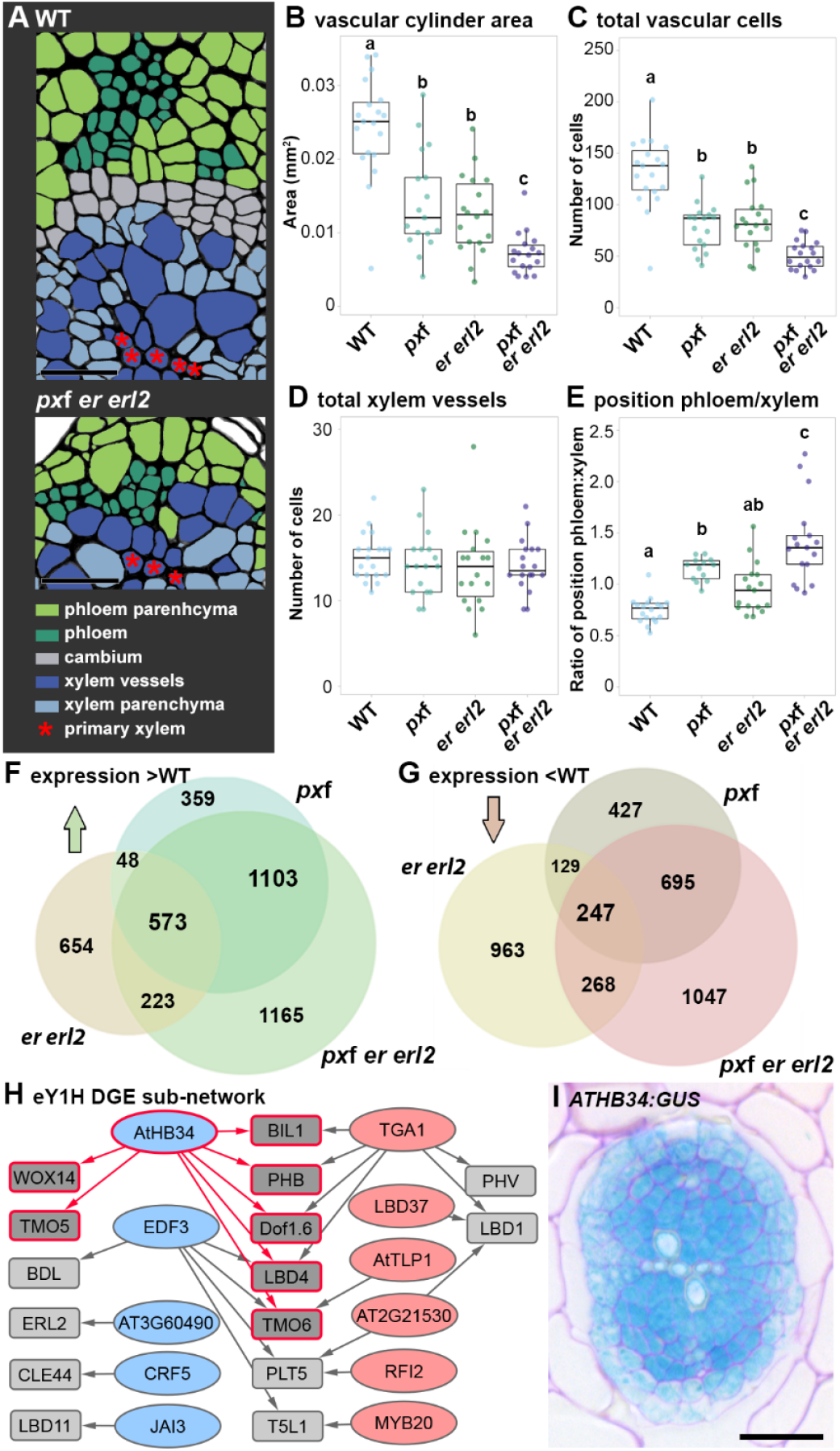
Homeodomain zinc finger transcription factors as candidate regulators of cambium development. (**A**) Diagrammatic representation of wild type (WT) and *px*f *er erl2* mutant 3-week-old roots. Cambium cells are not apparent in the *px*f *er erl2* line. Vascular area (**B**), vascular cell number (**C**), xylem vessel number (**D**), and ratio of position phloem:xylem relative to vascular cylinder centre (**E**; see also Supplementary Fig. S1A), in *px*f, *er erl2, px*f *er erl2* mutant plants and WT controls. (**F-G**) Venn diagrams showing overlap of differentially expressed genes between WT and *px*f; WT and *er erl2*; and WT and *er erl2 px*f lines. Values represent the number of genes significantly upregulated (F) or downregulated (G). (**H**) Transcriptional regulatory sub-network, containing prey proteins (ovals) in eY1H that were also differentially expressed in all three mutant backgrounds tested relative to WT. Rectangular nodes are bait promoters. Edges between nodes depict protein-DNA interactions. Up- and downregulated transcription factors in *px*f *er erl2* secondary growth regions are represented in red and blue, respectively. Bait nodes with red outlines highlight putative ATHB34 targets. (**I**) Expression pattern of *ATHB34:GUS* in 10-day-old roots. Scale bars are 20 μm (A) or 50 μm (I). Letters above boxplots (B-E) show homogeneous subsets (ANOVA plus Tukey post-hoc test).

Upon characterisation of changes in tissue composition, changes in gene expression in roots and hypocotyls, tissues undergoing active secondary growth, of *px*f, *er erl2*, and *px*f *er erl2* lines were determined using RNA-seq. Loss of *px*f, *er erl2*, or *px*f *er erl2* had profound effects on root mRNA profiles, with greater than 1000 genes up- or downregulated in the each of the mutant lines relative to wild type (adjusted *p* value ≤ 0.05; **Fig. 1f,g and Supplementary Data 1a-c**). *ER* was among the upregulated genes in *px*f lines, while *ERL1* and *PXY* were upregulated in *er erl2* roots (**Supplementary Dataset S1a,b**). This corroborates previous expression analyses (Wang et al. 2019), providing evidence for the reproducibility of the transcriptomes generated here. To maximise the likelihood of identifying regulators of cambium homeostasis, genes differentially expressed in all three mutants relative to wild type were identified. 247 genes were downregulated in the three mutant lines and 573 upregulated (adjusted *p* value ≤ 0.05), giving a combined total of 820 candidate genes from RNA-seq experiments (**Fig 1f,g** and **Supplementary Dataset S1d-e**).

To identify putative regulatory factors, we determined which of these 820 differentially expressed genes were able to bind to promoters of regulators of vascular development in enhanced yeast one-hybrid (eY1H) assays. This included a previously described network in which the promoters of 33 regulators of vascular development, including PXY and ER, had been screened against a library of vascular cylinder-expressed transcription factors yielding a network of 690 interactions (Gaudinier et al. 2011; Smit et al. 2020). An additional 744 transcription factor-promoter interactions were identified by screening promoters of a further six regulators of vascular development, *PLT5, LBD1, LBD11, WOX4, KNAT1*, and *DOF1*.*6*, against a comprehensive *Arabidopsis* transcription factor collection (Tang et al. 2021). This combined transcriptional regulatory network included a total of 1446 putative interactions between 38 promoters and 590 transcription factors (**Supplementary Fig. S1E and Supplementary Dataset S2**). Of the 820 candidate genes from the RNA-seq experiments in *px*f, *er erl2*, and *px*f *er erl2* lines, 11 formed a sub-network with expression of six transcription factors being upregulated and five downregulated in the absence of *PXY* family, *ER* and *ERL2* genes (**Fig. 1H**; blue nodes show downregulation, red show upregulation).

We focussed on ATHB34 as it was predicted to bind seven target promoters in the eY1H, making it the node with the greatest number of targets in the sub-network (**Fig. 1H**).

*ATHB34* encodes a plant-specific zinc finger homeodomain transcription factor (PLINC) of 255 amino acids (Tan and Irish 2006; Windhövel et al. 2001). In the context of the entire transcriptional regulatory network, *ATHB34* falls within a minority of highly connected proteins, as only 28 of the tested transcription factors (5%) occupied an excess of six promoters. The promoters to which ATHB34 bound in eY1H were those of *TMO5, BIL1, DOF1*.*6, PHB, WOX14, TMO6*, and *LBD4*. Given that *ATHB34* was identified in part due to its differential expression in lines in which secondary growth was perturbed (**Fig. 1B-G**), and that a subset of its putative targets in eY1H also regulate cambium activity (Etchells et al. 2013; Smet et al. 2019; Miyashima et al. 2019; Schlereth et al. 2010; De Rybel et al. 2014; Smit et al. 2020; Ye et al. 2021; Han et al. 2018; Kondo et al. 2014; Müller et al. 2016), we hypothesised that ATHB34 contributes to secondary growth regulation. In support, an *ATHB34:GUS* transcriptional reporter revealed promoter activity in the vascular cylinder of 10-day-old roots, suggesting *ATHB34* is expressed in secondary growth forming tissues (**Fig. 1I**).

### ATHB34 regulates the vasculature

To determine whether ATHB34 can influence vascular tissue patterns or proliferation, constitutive and inducible *ATHB34* over-expression lines were generated. In estradiol-induced *35S:XVE*>>*ATHB34* or constitutive *35S:ATHB34* roots at 10 days, the number of secondary xylem vessels was found to be greater than in wild type (**Fig. 2a,b and Supplementary Fig. S2**). Furthermore, in *35S:ATHB34* lines ectopic formation of secondary xylem cells was apparent, as determined by their presence in greater proximity to the epidermis than in controls (**Fig. 2c**). Thus, ATHB34 promoted differentiation of secondary xylem vessels.

**Figure 2:**
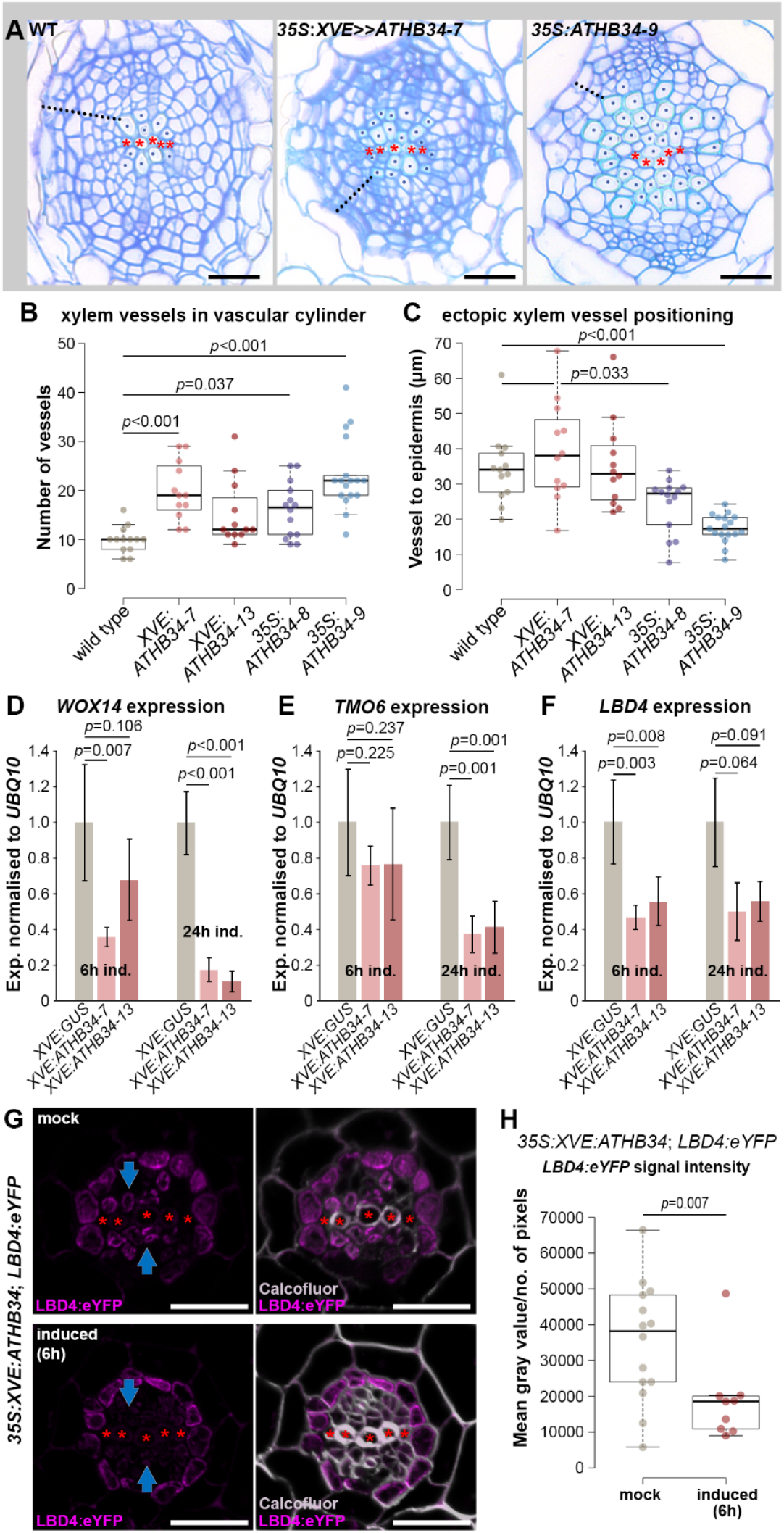
ATHB34 promotes xylem differentiation and represses the WOX14-TMO6-LBD4 feedforward loop. (**A-C**) Transverse section through wild type (WT), induced *35S:XVE*>>*ATHB34-7*, and *35S:ATHB34-9* roots. Red asterisks define the primary xylem. Secondary xylem vessels, quantified in (**B**) are marked with blue dots. Dashed line shows shortest distance from xylem vessel to periderm, quantified in (**C**). (**D-F**) qRT-PCR analysis showing reduced expression of *WOX14, TMO6*, and *LBD4* in *35S:XVE:ATHB34* lines compared to an *XVE:GUS* control upon 6 h and 24 h estradiol induction. (**G-H**) The effect of *35S:XVE:ATHB34* activation on an *LBD4:eYFP* reporter in root transverse sections. *LBD4:eYFP* signal is lost from the cambium (blue arrows) upon induction (**G**), as reflected by measurements of signal intensity (**H**). *p* values were calculated using ANOVA plus Tukey (B, C) or Dunnett (D-F) post-hoc test. A Student’s t-test was used in (H). Scale bars (A, G) are 20 μm.

Next, we sought to address how ATHB34 was regulating differentiation. We have previously shown that a coherent feedforward loop comprised of *WOX14, TMO6*, and *LBD4* promotes vascular cell division (Smit et al. 2020). ATHB34 bound to the promoters of these three genes in eY1H, leading to the hypothesis that ATHB34 might repress their expression to promote differentiation. Indeed, 6 h- or 24 h-induced *35S:XVE*>>*ATHB34* seedlings demonstrated reduced *WOX14, TMO6*, and *LBD4* expression in comparison to a *35S:XVE*>>*GUS* control, determined by qRT-PCR (**Fig. 2d-f**). We then focussed on *LBD4* as it is the output gene of the WOX14-TMO6-LBD4 feedforward loop(Smit et al. 2020). An *LBD4:eYFP* transcriptional reporter was transformed into *35S:XVE*>>*ATHB34* plants and accumulation of eYFP was determined in root transverse sections in the presence or absence of a 6 h estradiol induction. In mock treatments, *LBD4:eYFP* expression was evident in the cambium and pericycle cells as described previously(Ye et al. 2021) but was absent from the cambium of estradioltreated *35S:XVE*>>*ATHB34 LBD4:eYFP* plants (**Fig. 2G**). After quantifying eYFP signal intensity across the vascular tissues, a reduction was observed upon induction of *ATHB34* (**Fig. 2H**).

Having demonstrated that ATHB34 is a negative regulator of *WOX14, TMO6*, and *LBD4* expression, we aimed to provide further confirmation that ATHB34 is sufficient to bind to the promoters of these three genes *in planta*. The promoters of *WOX14, TMO6*, and *LBD4* were fused to a minimal *35S* promoter (*m35S*) upstream of a *LUC* reporter, in a plasmid that also carried an estradiol-inducible *ATHB34* cassette. When the *LBD4pro:m35S:LUC*-*35S:XVE*>>*ATHB34* plasmid was transfected into *Nicotiana* leaves, the LUC signal was greater when leaves where treated with mock treatment than with estradiol (*ATHB34* induction) (**Fig. 3A**). This supports the idea that ATHB34 binds to the *LBD4* promoter to repress gene expression *in planta*. Similar observations were observed for the *TMO6* promoter, the *WOX14* promoter, and the *TMO6* intron (**Fig. 3A-E**) which was included due to evidence of ATHB34 binding in DAP-seq data (**Fig. 3F**)(O’Malley et al. 2016).

**Figure 3.**
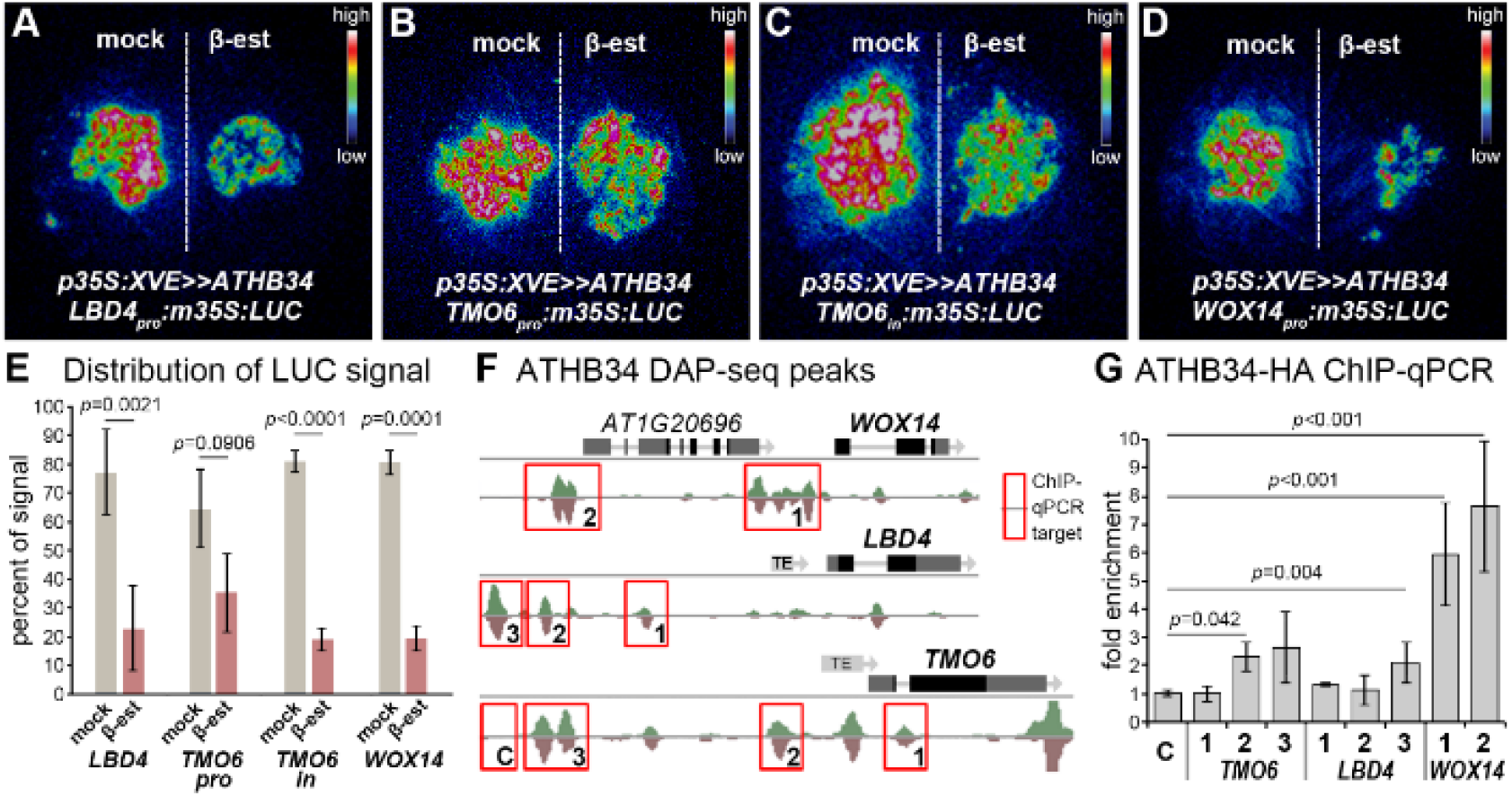
ATHB34 is a direct repressor of *WOX14, TMO6*, and *LBD4* transcription. (**A**-**E**) Effects of *35S:XVE*>>*ATHB34* on an *LBD4pro:m35S:LUC* (**A**), *TMO6pro:m35S:LUC* (**B**), *TMO6in:m35S:LUC* (**C**), or *WOX14pro:m35S:LUC* (**D**) reporter in *Nicotiana* leaves where the left side of leaves was mock treated and the right induced. The percentage of the total LUC signal per leaf on the mock and treated sides of the leaf is shown in (**E**). (**F**) Diagram showing ATHB34 DAP-seq binding peaks in the *WOX14, TMO6*, and *LBD4* genic region^30^ with red boxes marking areas subsequently targeted using ChIP-qPCR (**G**). (**G**) CHIP-qPCR testing ATHB34-HA occupancy of *WOX14, TMO6*, and *LBD4* promoters and the *TMO6* intron in *Arabidopsis*. X-axis labels correspond to primer locations within red boxes in (F). *p* values in G were calculated using ANOVA plus Dunnett tests with those less than 0.05 relative to the control marked. In (E) all *p* values are shown, determined using Students’ t-tests.

ATHB34 DAP-seq data(O’Malley et al. 2016) was also consulted to identify regions within the *WOX14, TMO6*, and *LBD4* promoters that could be tested for ATHB34 occupancy *in planta* using ChIP-qPCR. Having identified putative binding regions (**Fig. 3F**), *Arabidopsis* plants harbouring an *ATHB34pro:ATHB34-HA* fusion were subjected to ChIP-qPCR. DNA associated with immunoprecipitated ATHB34 was enriched in a subset of *WOX14, TMO6* and *LBD4* promoter regions (**Fig. 3G**). As such, ATHB34 bound to the promoters of *WOX14, TMO6*, and *LBD4 in vitro* (DAP-seq(O’Malley et al. 2016)), in yeast (eY1H; **Fig. 1H**), in *Nicotiana* (LUC assay; **Fig. 3A-E**), and *in vivo* (ChIP-qPCR; **Fig. 3G**), demonstrating that ATHB34-mediated regulation of these genes occurs via direct interaction with their promoters.

ATHB34 redundancy with ATHB30 and ATHB23 Interrogation of publicly accessible Arabidopsis co-expression data(Obayashi et al. 2018) identified two related genes, *ATHB23* and *ATHB30* with similar expression profiles to *ATHB34* (**Supplementary Dataset S3**). A maximum likelihood phylogeny constructed based on alignment of

PLINC family protein sequences suggested that ATHB34, ATHB23 and ATHB30 form a well-supported clade (**Fig. 4a**). Furthermore, *ATHB23* and *ATHB30* expression was investigated within our RNA-seq dataset. By contrast to *ATHB34*, which demonstrated expression reductions in *px*f, *er erl2*, and *px*f *er erl2* roots, the expression of both *ATHB23* and *ATHB30* was elevated in *px*f *er erl2. ATHB23* expression was also upregulated in *px*f (adjusted *p* values < 0.05; **Fig. 4b**). Genes frequently exhibit a compensatory increase in expression when that of a redundantly acting homologue is attenuated(Kafri et al. 2009). We thus hypothesised that these genes function redundantly.

**Figure 4.**
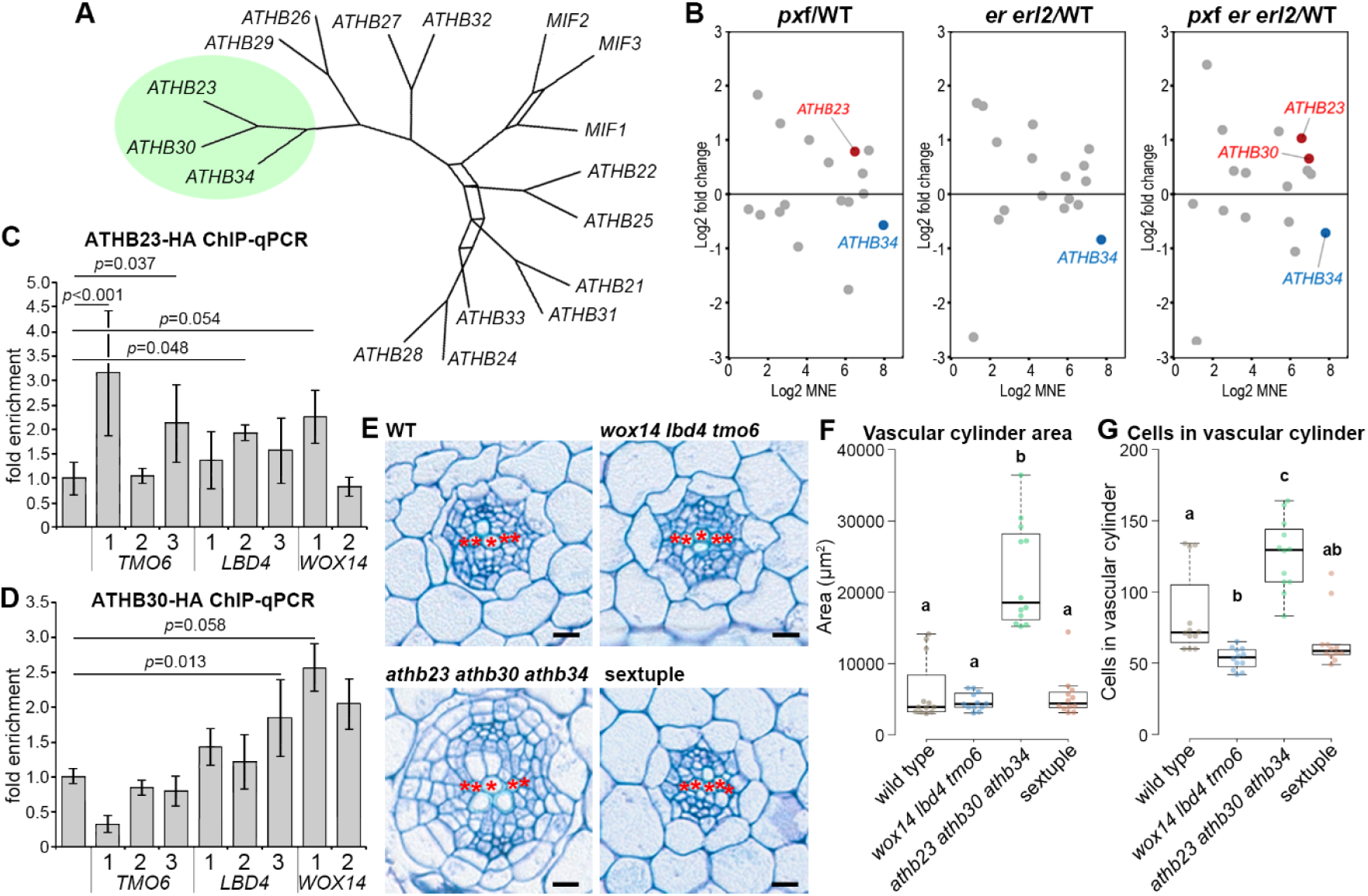
ATHB34 acts redundantly with ATHB30 and ATHB23. (**A**) Consensus network showing support for the grouping of PLINC family protein sequences, based on a maximum likelihood tree and bootstrapping analysis with 100 replicates. (**B**) Mean fold change in expression and mean normalised expression (MNE) of all members of the PLINC transcription factor family in *pxf, er erl2* and *pxf er erl2* hypocotyls compared to wild type at 3 weeks old. *ATHB23, ATHB30*, and *ATHB34* are shown in red and blue where significantly up- and downregulated (Wald test, BH adjusted *p* ≤ 0.05, |log2 fold change| ≥ 0.0). (**C**-**D**) ChIP-qPCR showing the binding of ATHB23 (**C**) and ATHB30 (**B**) to the promoters of *WOX14, TMO6*, and *LBD4*, and the *TMO6* intron in the regions marked in Figure 3F. (**E-G**) *athb23 athb30 athb34* triple mutants at 16 days old, showing increased radial growth in comparison to wild type. This increase is attenuated in *athb23 athb30 athb34 wox14 lbd4 tmo6* sextuple mutants, shown in transverse sections (**E**), vascular cylinder area measurements (**F**) and vascular cell numbers (**G**). *p* values were calculated using an ANOVA with a Dunnett post-hoc test (C-D), or Tukey post-hoc test where letters above the boxes show homogeneous subsets (F-G). Red asterisks define the primary xylem, and scale bars are 20 μm (E).

To further test this idea, we determined whether, in addition to ATHB34, ATHB23 and ATHB30 interacted with *WOX14, TMO6*, and *LBD4* regulatory DNA. Within the eY1H network ATHB23 and ATHB30 demonstrated similar levels of connectedness to ATHB34 (**Supplementary Fig. S3a**). Upon visualisation of a sub-network containing the PLINC transcription factors, ATHB23 was found to bind the *WOX14* and *LBD4* promoters, while ATHB30 bound those of *WOX14, TMO6*, and *LBD4* (**Supplementary Fig. S3b**). Plasmids carrying *WOX14pro:m35S:LUC, TMO6pro:m35S:LUC, TMO6intron:m35S:LUC*, and *LBD4pro:m35S:LUC* were again infiltrated into *Nicotiana* leaves, in combination with an estradiol-inducible *ATHB23* or *ATHB30* cassette and either an estradiol or a mock treatment. As was the case with *ATHB34* (**Fig. 3a-d**), induction of either *ATHB23* or *ATHB30* resulted in a reduced luciferase signal (**Supplementary Fig. S3c,d**). ChIP-qPCR was also performed using *ATHB23pro:ATHB23-HA* or *ATHB30pro:ATHB30-HA* plant lines. Here, nuances were observed. ATHB23 demonstrated binding to the *TMO6* intron, which was not observed for either ATHB30 or ATHB34. ATHB23 also bound to the *LBD4* promoter, as did ATHB30 (**Fig. 4c,d and Fig. 3g**). As such, regulatory regions of feed-forward loop genes were occupied by ATHB23 and ATHB30 in addition to ATHB34, supporting the idea that these three proteins act redundantly.

Taking a loss-of-function genetic approach to further explore roles in secondary growth, *athb23, athb30*, and *athb34* T-DNA insertion mutants were identified, and triple and double mutants generated by crossing. *athb23 athb30 athb34* triple mutants exhibited increased radial growth relative to the wild type or single mutants, determined by analysing root transverse sections. Indeed, the area of the root vasculature in transverse section increased concomitantly with the number of root vascular cells (**Supplementary Fig. S4)**. ATHB34 appears to be a critical component as any double or triple mutant lacking ATHB34 demonstrated the growth increase, while *athb23 athb30* doubles were indistinguishable from wild type. Because ATHB34 represses the expression of cambium promoting factors (**Fig. 2d-h**), in *athb23 athb30 athb34* lines, cells leaving the cambium that would otherwise differentiate in wild type, likely retain meristem identity and mitotic competence for longer. This, in turn would lead to increased cell production. We tested whether the increase in cell division observed in *athb23 athb30 athb34* depended on the presence of *WOX14, TMO6*, and *LBD4*. An *athb23 athb30 athb34 wox14 tmo6 lbd4* sextuple mutant was generated by transforming *wox14 tmo6 lbd4* lines with a genome editing construct carrying guide RNAs targeting *ATHB34, ATHB30*, and *ATHB23*. The root vascular cylinder area and the number of cells in vascular cylinder transverse sections of sextuple mutant plants was similar to those of wild type and *wox14 tmo6 lbd4* lines, and smaller than that of *athb23 athb30 athb34* lines (**Fig. 4e-g**). Thus, the *athb23 athb30 athb34* enhanced secondary growth phenotype was suppressed in the *wox14 tmo6 lbd4* background.

## Discussion

Secondary growth depends on cell division within the cambium, with the major positive regulators of cambium activity being TDIF-PXY signalling, auxin, and cytokinin, which in turn promote expression of transcription factors WOX14, TMO6, and LBD4, among others (Smit et al. 2020; Smet et al. 2019; Ye et al. 2021; Etchells et al. 2013). However, meristems typically act in balance, that is, negative regulators are required to dampen activity. Here, negative regulators were identified by cross-referencing transcriptome data against an eY1H transcriptional network. The network constituted a previously described set of protein-DNA interactions centred on PXY signalling, supplemented by new screens to identify proteins acting upstream of central regulators of cambium function or meristem activity, DOF1.6, WOX4, LBD1, LBD11, PLT5 and KNAT1. This expanded eY1H network provided a resource for hypothesis building, leading to identification of ATHB34 and its homologues, ATHB23 and ATHB30 as repressors of cambium activity. The finding that these proteins bind directly to *WOX14, TMO6* and *LBD4* promoters, leading to attenuation of transcription, provides a mechanistic framework for balancing cambium activity.

Questions remain regarding ATHB34, ATHB23, and ATHB30 function. Firstly, upstream regulators are not described. However, like ATHB34, ABA is known to promote xylem differentiation (Campbell et al. 2018; Ramachandran et al. 2021) and in publicly available datasets, *ATHB34* and *ATHB30* expression is activated by ABA (Okamoto et al. 2012; Böhmer and Schroeder 2011), providing a hypothesis for further investigation. Secondly, a question arises from the observation that ATHB34 repression of *LBD4* expression was not uniform across the root. *LBD4* is expressed in the cambium and the periderm, but ubiquitous activation of ATHB34 attenuated *LBD4* expression specifically in the cambium and not the periderm (**Fig. 2G**). This suggests a vascular tissue-specific element of ATHB34 activity remains to be discovered. Possibilities include a vascular cylinder-specific protein binding partner such as BES1 or a HD-Zip III transcription factor, or a tissue-specific protein modification. Identification of this unknown component is a matter for further research. *ATHB34* expression is broad and as such, a spatially restricted component that protects the cambium from its activity must be a component of the system.

In summary, we previously identified a series of transcription factors that promote cambium activity and stem cell fate. Here we have identified a family of zinc finger homeodomain transcription factors that oppose that activity.

## Acknowledgements

CC gratefully acknowledges the National Key Research and Development Program of China (2024YFE0102300), the National Natural Science Foundation of China (32470578), Advanced Foreign Experts Project (H20250816), and the Fundamental Research Funds for the Central University (2662024SZ001) for funding support. XW gratefully acknowledges the National Natural Science Foundation of China Youth Fund (32400098). XW, QH and JPE thank the Chinese Scholarship Council (CSC) Scholarship support for visiting Durham University and doctoral training grants, respectively. JPE, APM, JH, RB, and QH thank UKRI (BBSRC grants BB/V008129/1 and UKRI2966), and APM and RM thank the European Research Council (ERC-CoG CORKtheCAMBIA, agreement no. 819422) for funding. AMB and SMB were funded by an HHMI Faculty Scholar award and the UC Davis Genome Center.

## Materials and methods

### Accession numbers

Arabidopsis Genome Initiative (AGI) numbers and descriptions of all genes identified by RNA-seq are provided in Supplementary Data 1A-F. RNA-seq data is available on GEO (accession number GSE316512). AGI numbers corresponding to Arabidopsis promoters and transcription factors comprising the eY1H networks utilised by this research are provided in Supplementary Data 2.

### Plant materials

The Arabidopsis ecotype Columbia-0 (Col-0, referred to as wild type) and previously described *pxy-3 pxl1-1 pxl2-1* triple mutant (*px*f)(Fisher and Turner 2007), *er-105 erl2-1* double mutant (*er erl2*) and *px*f *er erl2* quintuple mutant lines(Wang et al. 2019; Shpak et al. 2005), were obtained from laboratory stocks. Homozygous T-DNA insertion lines, *athb34-1* (SALK_ 085482C), *athb30-1* (GABI-Kat_ 363F04) and *athb23-1* (SALK_ 041382C), were obtained from the Nottingham Arabidopsis Stock Centre (NASC). Triple and double mutants were generated by crossing. To obtain sextuple *athb23 athb30 athb34 wox14 tmo6 lbd4* lines, a genome editing construct carrying guide RNAs against *ATHB23, ATHB30* and *ATHB34* was introduced to *wox14 tmo6 lbd4* triple mutants(Smit et al. 2020). Sextuple mutants were selected by sequencing. *ATHB34:GUS* was generated by cloning the *ATHB34* promoter into pGWB3(Nakagawa et al. 2007) via pENTR-D-TOPO. *35S:ATHB34, 35S:XVE:ATHB34* were generated similarly using the *ATHB34* CDS and destination vectors pMDC7(Curtis and Grossniklaus 2003) and pK2GW7(Karimi et al. 2002). *ATHB34pro:ATHB34-HA, ATHB30pro:ATHB30-HA, ATHB23pro:ATHB*23*-HA* were assembled using Greengate cloning(Lampropoulos et al. 2013). Vectors were transformed into Arabidopsis plants using floral dipping (Clough and Bent 1998). Homozygous lines were selected in the T3 generation.

### Imaging

For phenotyping, thin sections taken from plants embedded in JB4 resin (Polysciences) were generated. The *ATHB34:GUS* reporter was stained, embedded in Technovit 7100 (Kulzer), sectioned and counterstained with ruthenium red prior to imaging. Sections were mounted with Histomount (National Diagnostics) and visualized on a Zeiss Axioskop microscope. Confocal imaging was performed on fresh material which had been embedded in 4% agarose prior to sectioning on a 7000smz-2 vibratome (Campden Instruments). The sections were incubated in calcofluor prior to visualisation on a Zeiss 880 confocal microscope.

### Transcriptome analysis

Plants were grown under standard long day conditions. Total RNA was extracted from hypocotyls and the upper 5 mm of roots of 3-week-old wild type, *px*f, *er erl2* and *px*f *er erl2* plants. Paired-end RNA-seq, including library preparation and sequencing using Illumina NovaSeq 6000, were conducted by Novogene. Sequencing reads were mapped to the Arabidopsis TAIR10 genome (EnsemblePlants, release 47) with the corresponding gtf file using HISAT2 (v2.0.5)(Kim et al. 2019). The read numbers mapped to each gene were counted using featurecounts (v1.5.0-p3)(Liao et al. 2014). Differential expression analysis and normalisation of read counts using the median of ratios method were conducted in R (v3.6.2) using DESeq2(v1.20.0)(Love et al. 2014). For each mutant genotype, log2 fold change of mean normalised counts in mutant (n = 3) vs wild type (n = 3) samples were calculated, and Wald tests conducted under the null hypothesis that genes were equally expressed across genotypes. The resulting *p* values for each gene were adjusted using the Benjamini-Hochberg (BH) procedure to correct for multiple testing. Analysis of the overlap of significantly differentially expressed gene sets (Wald test, BH adjusted p ≤ 0.05, |log2 fold change| ≥ 0.0) was carried out using BioVenn(Hulsen et al. 2008).

For co-expression analysis the ATTED-II v10.1 co-expression database was used to access publicly available Arabidopsis microarray and RNA-Seq data, and calculate mutual ranks based on linear regression of available datasets(Obayashi et al. 2018).

### Transcriptional regulatory networks

Physical interactions between Arabidopsis transcription factors and promoters of known regulators of vascular development were predicted by eY1H assays. Screens against *DOF1*.*6, WOX4, LBD1, LBD11, PLT5* and *KNAT1* were performed as previously described(Gaudinier et al. 2011; Reece-Hoyes et al. 2011; Gaudinier et al. 2017), using a transcription factor collection previously described(Tang et al. 2021). The gene regulatory network was generated by combining the data generated here with one previously described for TDIF-PXY signalling(Smit et al. 2020) (Supplementary Data 2). Networks and subnetworks, were visualised using Cytoscape v3.8.2(Shannon et al. 2003).

### Phylogenetic analysis

Amino acid sequences for 17 Arabidopsis PLINC family genes were retrieved from NCBI and aligned using PROMALS3D(Pei et al. 2008). A maximum likelihood phylogeny was constructed based on a JTT model of protein evolution using the R package, *phangorn*(Schliep 2011). Bootstrapping support values were calculated based on 100 replicates. A consensus network was constructed to visually represent the support for each branch, with longer branch lengths indicating greater support.

### qRT-PCR analysis

Seedlings were collected at the designed time points and immediately frozen in liquid nitrogen. mRNA extraction and cDNA synthesis were performed using Sera-Mag Oligo-dT beads. qRT-PCR was conducted according to the steps described in a qPCRBIO SyGreen Mix Lo-ROX (PCRBiosystems, UK) protocol, and qPCR reactions were run in a CFX Duet Real Time system (Bio-Rad). The amplified *UBQ10* transcript abundance was used as a normalisation control, and relative expression values were calculated using the 2^-ΔΔCT^ method. qPCR was performed with three biological and three technical replicates. All possible pairwise comparisons were tested using a one-way ANOVA followed by a post-hoc Tukey HSD test.

### Luciferase reporter assay

To confirm protein-DNA interactions *in vivo*, a luciferase reporter assay was performed. The promoters of *WOX14, LBD4*, and *TMO6*, and the *TMO6* intron were fused upstream of a constitutively transcribed *LUCIFERASE (m35S:LUC)* reporter. These cassettes were then subcloned into pER8, a vector also conferring estradiol-inducible expression with the CDS of *ATHB23, ATHB30*, and *ATHB34* each cloned under the control of the *35S:XVE* promoter. This generated a vector in which an *ATHB* transcription factor was inducible, and the promoter to which it putatively bound was controlling luciferase expression within the same plasmid. The constructs were transformed into *Agrobacterium* GV3101 and then infiltrated into *Nicotiana benthamiana* leaves. The transformed plants were placed in the dark for 24 h and then under a 12 h day/night cycle for 48 h. At 48 h after infiltration, leaves were treated with either mock solution or β-estradiol. Luciferase activity was analysed 24 h after treatment by spraying leaves with 1 mM luciferin and detecting luminescence using a chemiluminescence imaging system.

### ChIP-qPCR analysis

8-day-old *ATHB23pro:ATHB23:HA, ATHB30pro:ATHB30:HA*, and *ATHB34pro:ATHB34:HA* seedlings were collected for chromatin immunoprecipitation (ChIP) assays, which were performed as previously described (Ghosh et al. 2022). Samples were cross-linked in 1% (w/v) formaldehyde and then ground into powder using liquid nitrogen. The crude nuclear pellet was resuspended in nuclear lysis buffer and sonicated at 6 °C for 6 min using a Covaris M220 focused-ultrasonicator. 1 μg of Pierce anti-HA magnetic beads (Thermo) was added to the soluble chromatin solution and incubated overnight for immunoprecipitation of protein-DNA complexes. The isolated chromatin before precipitation was used as an input control. 10% Chelex slurry (Bio-Rad) was used for the de-crosslinking reaction. To identify the genomic regions regulated by ATHB23, ATHB30, and ATHB34, primer pairs were designed to amplify four distinct regions within the promoters of *WOX14, TMO6*, and *LBD4*. qPCR reactions were run in a CFX Duet Real Time system (Bio-Rad). ChIP-qPCRs were performed with three biological and three technical replicates. Experimental samples were compared to controls using an ANOVA, followed by a post-hoc Dunnett test.

## Supplemental figures and legends

**Figure S1.**
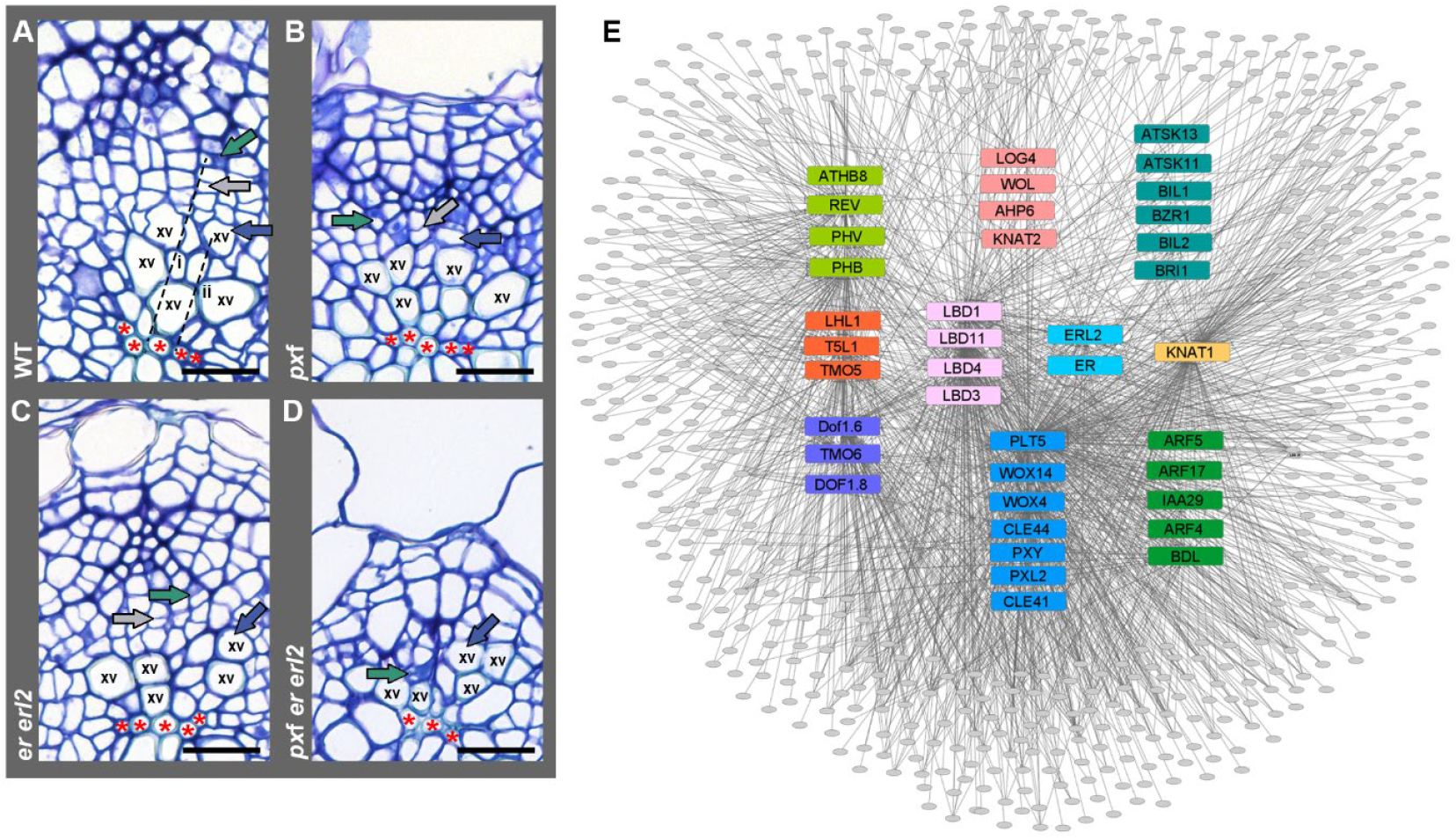
Mediators of secondary growth. (**A-D**) Transverse sections from *px*f *er1 erl2* mutants and controls. (**A**) Wild-type (WT), (**B**) *er*f, (**C**) *er erl2*, (**D**) *px*f *er erl2 exf* vascular bundles. Xylem, cambium and phloem are marked with blue, grey and green arrows, respectively. Red asterisks define the primary xylem. In (A), dashed line (i) marks the distance from root centre to phloem, while line (ii) marks the distance from the primary xylem at the root centre to differentiating xylem. The ratio of ii:i is graphed in Figure 1E. xv denotes secondary xylem vessels. Scales bars are 20 μm. (**E**) Schematic representation of the expanded transcriptional regulatory network. Promoters are represented by coloured rectangles, and transcription factors by ovals. Edges between nodes depict protein-DNA interactions.

**Figure S2.**
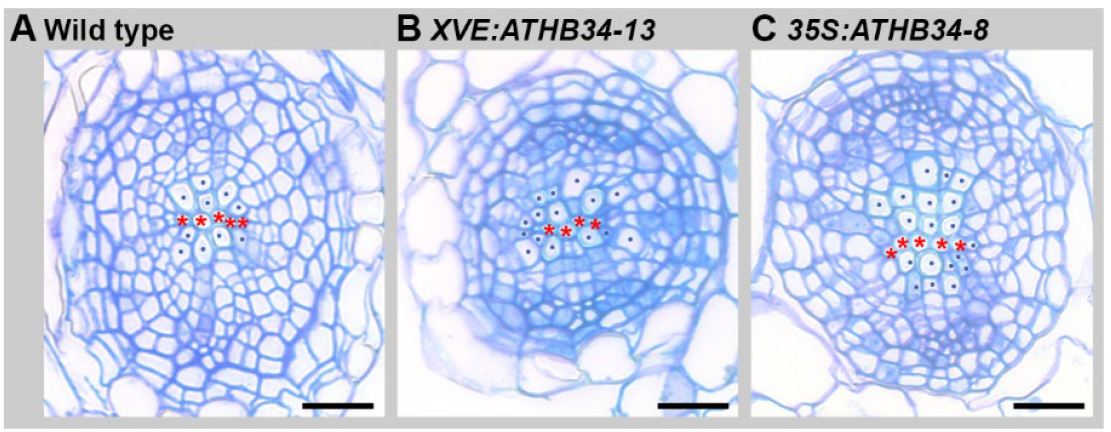
Phenotypes of vascular tissue in *ATHB34* overexpression lines. Transverse section through wild type (**A**), induced *35S:XVE*>>*ATHB34-13* (**B**), and *35S:ATHB34-8* (**C**) roots. Red asterisks define the primary xylem. Secondary xylem vessels are marked with blue dots. Scale bars are 20 μm.

**Figure S3.**
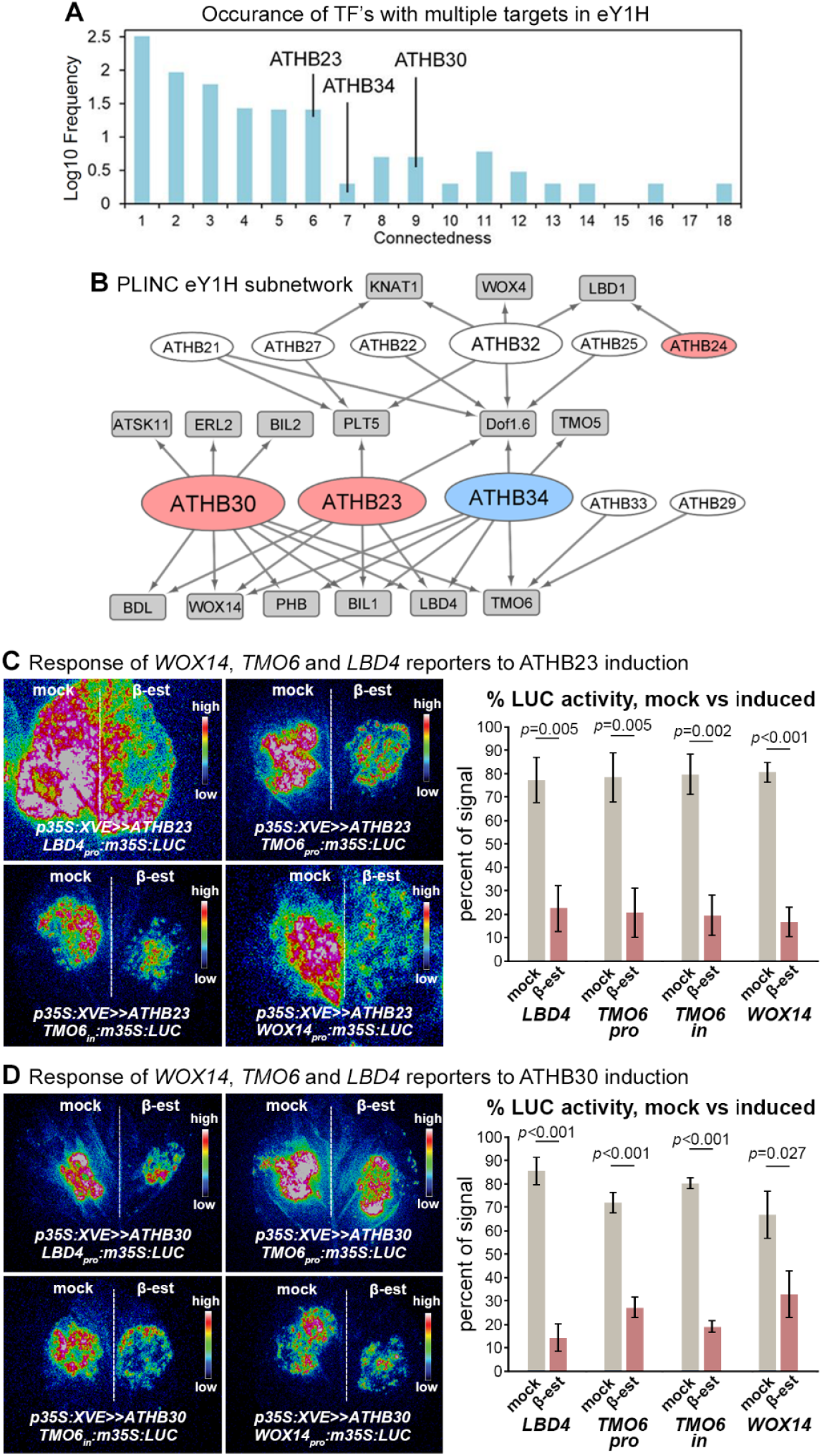
ATHB23 and ATHB30 transcriptional network. (**A**) Distribution of transcription factors (TF’s) that bind to multiple targets in eY1H. The connectedness of ATHB34, ATHB23, and ATHB30 are marked. (**B**) Transcriptional regulatory sub-network containing PLINC transcription factors. Up- and downregulated transcription factors in *pxf er erl2* hypocotyls are represented in red and blue, respectively. Edges between nodes depict protein-DNA interactions. Node size represents connectedness. (**C-D**) Effects of *35S:XVE*>>*ATHB30* or *35S:XVE*>>*ATHB23* on *LBD4pro:m35S:LUC, TMO6pro:m35S:LUC, TMO6in:m35S:LUC*, and *WOX14pro:m35S:LUC* reporters in *Nicotiana* leaves. The left side of leaves were mock-treated and the right induced. The percentage of the total signal on each side of leaves is shown in the histograms. *p* values were determined using Student’s t-tests.

**Figure S4.**
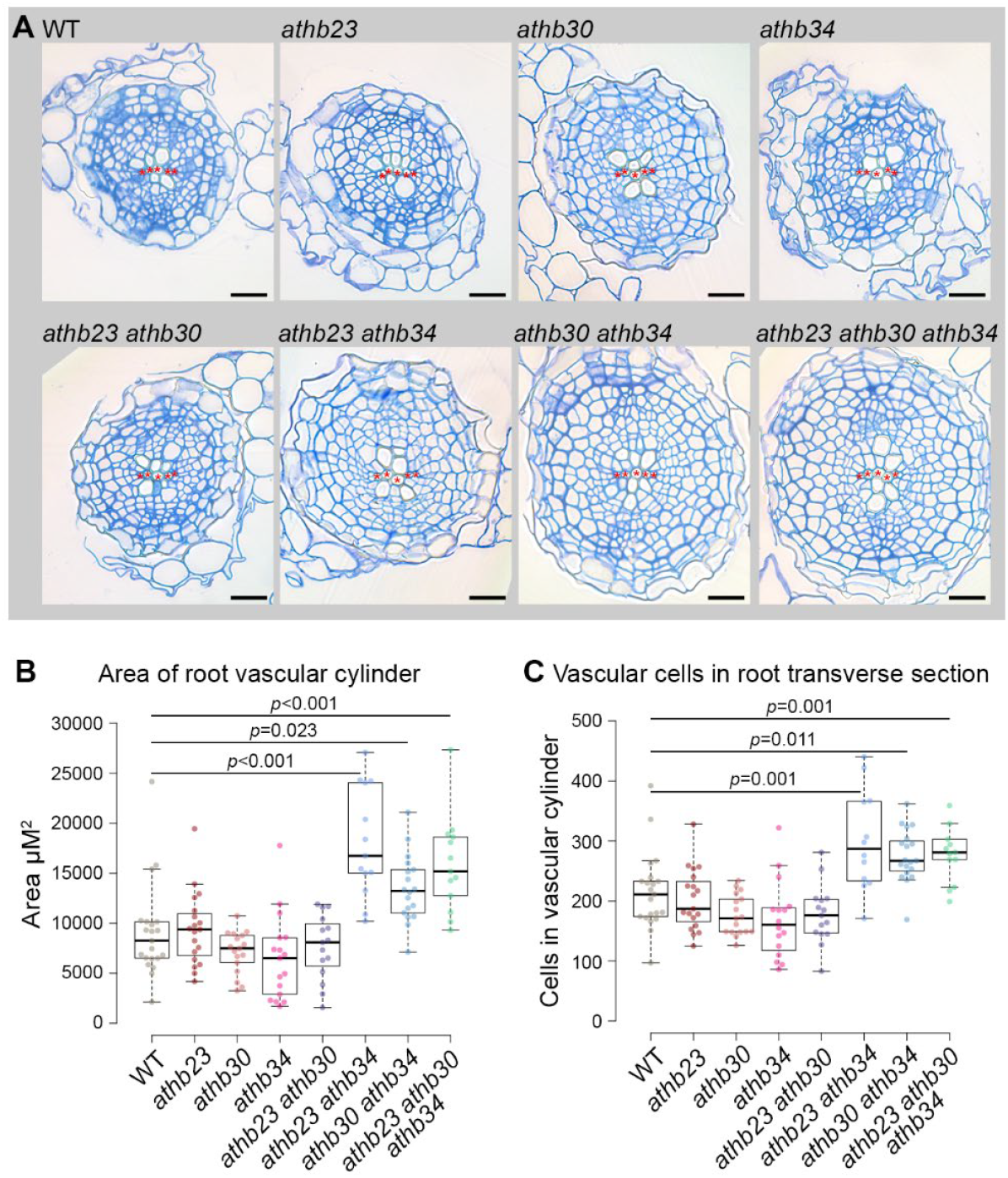
Comparison of *athb23 athb30 athb34* lines and controls. (**A**) Wild-type (WT) and *athb23 athb30 athb34* triple mutants alongside single and double mutant controls. (**B-C**) Boxplots show the vascular cylinder area (**B**) and vascular cell numbers (**C**). *p* values were calculated using an ANOVA with a Tukey post-hoc test. Significant differences are marked. Red asterisks define the primary xylem. Scales bars are 20 μm.

